# Antimicrobial Resistance Profiling and Phenotypic Characterization of Archived Clinical *Bacillus paranthracis* Strains

**DOI:** 10.64898/2026.04.16.719033

**Authors:** Pierre Michel, Tucker Maxson, Vasanta Chivukula, Will Overholt, Luz Karime Medina Cordoba, Stephanie Ayodele-Abiola, John McQuiston, Cari A. Beesley, Melissa Bell, Victoria Caban Figueroa, Julia Bugrysheva, Tyler Chandross-Cohen, Zachary Weiner, Laura Carroll, Jasna Kovac, David Sue

## Abstract

*Bacillus paranthracis* was formally defined as a species in 2017, after decades of carrying the name “emetic B. cereus” based on cereulide production and clustering within *B. cereus sensu lato phylogenetic* group III. Commonly associated with foodborne intoxication, reports rarely link *B. paranthracis* to non-foodborne clinical illness. As such, the new taxonomy and close resemblance of the name to the biothreat pathogen *Bacillus anthracis* cause confusion in diagnostic and public health settings. To address this issue, *B. paranthracis* clinical strains (n=20) from the CDC collection were tested with microbiological methods used for identification of *B. anthracis* and antimicrobial susceptibility testing. Some *B. paranthracis* phenotypes were similar to *B. anthracis,* however others were inconsistent across strains. Like *B. anthracis*: 3 strains tested capsule positive, 5 were non-hemolytic on blood agar, and 9 non-motile. All *B. paranthracis* strains were resistant to gamma phage lysis, which differentiated them from *B. anthracis*. Treatment regimens for *B. paranthracis* infections are not well established, as antimicrobial therapy is not indicated for emetic intoxication caused by *B. paranthracis*. Notably, six *B. paranthracis* strains had elevated minimal inhibitory concentrations to anthrax-recommended antibiotics: one for ciprofloxacin, three for doxycycline and tetracycline, and two for clindamycin. Rapid MinION sequencing was assessed for antimicrobial resistance detection prediction but had limited value when using PiMA v.1. These microbiological observations and susceptibility profiles of *B. paranthracis* expand our understanding of this pathogen, strengthening our ability to differentiate this bacterium from *B. anthracis* to improve diagnosis and patient outcomes.

**IMPORTANCE:** This study describes *in vitro* characterization of 20 archived clinical strains of *B. paranthracis*, an opportunistic pathogen identified more frequently in recent reports. Our findings highlight phenotypic differences and similarities between *B. paranthracis* and *B. anthracis* using standard microbiological methods and drug susceptibility profiling. We also assess a rapid *B. anthracis* specific MinION long read genome sequencing workflow with *B. paranthracis*. This report highlights the overlapping morphological features shared by *B. paranthracis* and *B. anthracis* to improve future laboratory diagnosis and strengthen anthrax preparedness. This article will effectively reach an audience of public health professionals and microbiologists strengthening anthrax preparedness.

## INTRODUCTION

*Bacillus paranthracis* (“like” or “near” *Bacillus anthracis*) was first named in 2017 based on type strain (called “Mn5”) isolated from marine sediment [1]. *B. paranthracis* was proposed as a novel species using 16S rRNA gene sequences, multilocus sequence typing analysis (MLST), digital DNA–DNA hybridization (dDDH) and average nucleotide identity (ANI) comparisons with other *Bacillus cereus sensu lato* (*B. cereus s.l.*), a species comprised of many closely related lineages [1, 2]. *B. paranthracis* was previously undifferentiated from *B. cereus* or “group III *B. cereus”* which include *B. cereus s.l.* strains that have the potential to cause emetic/diarrheal foodborne illnesses [3]. As of March 2026, less than 100 articles (NCBI query) reference *B. paranthracis* and few are associated with human illness. By contrast, the biothreat agent *B. anthracis* [4] (causative agent of anthrax), is referenced 7000+ times in NCBI PubMed.

*B. anthracis* is a globally pervasive, soil dwelling organism that causes severe illness and death in humans and other animals through direct contact [5]. The infections are serious and demand both timely and accurate diagnosis [6]. As a member of the *B. cereus s.l.* by taxonomy, *B. anthracis* has notably low genetic diversity across strains when compared to other group members within its species [7].

*B. cereus s.l.* bacteria sporulate, have low GC content, are environmentally pervasive, but can vary in other properties including virulence to humans [8]. Previous reports implicated *B. paranthracis* in foodborne and waterborne outbreaks, as a coinfection in a fatal Ebola virus disease case, and as an opportunistic pathogen [9-13]. Sabin, et al. described the genetic features of 20 clinical *B. paranthracis* strains in a genome study of 93 clinical *B. cereus s.l.* archival strains (1967-2003) and 5 clinical *B. cereus s.l*. contemporary strains (2019-2023) from the CDC collection[14]. Although *B. paranthracis* strains were previously reported as lacking the anthrax virulence genes found on *B. anthracis* plasmids pXO1 and pXO2 [1], which cause toxemia and facilitate immune system evasion through capsule formation [15], Sabin et. al identified anthrax-associated virulence genes (*capBCADE* operon) in three B. *paranthracis* strains [14].

The taxonomic distinctions between *B. anthracis* and *B. cereus s.l.* species are not always clear cut, due to genetic similarities, the presence of virulence genes/plasmids, and observations of phenotypic variation [2]. Nomenclature for some *B. cereus s.l*. species might also complicate this issue. To distinguish anthrax-toxin producing *B. cereus s.l.* strains, the term “*biovar Anthracis*” was added to the genus/species name[3, 16]. Notably, this highly similar naming scheme shared by species with known virulence, or lack thereof, has delayed outbreak investigations over nomenclature confusion, based on concerns over strains labeled as *B. paranthracis, B. anthracis,* and *B. cereus biovar anthracis* [3]. Accurate laboratory detection of *B. paranthracis* (and differentiation from *B. anthracis*) aids clinicians in diagnostic interpretation and decision-making, while de-escalating unnecessary alarm caused by the suspicion of human anthrax infections.

Human anthrax treatment guidelines were recently updated and include empiric treatment and post exposure prophylaxis guidance that cover different routes of infection (cutaneous, inhalation, gastrointestinal, injection) [17]. Clinical guidance for other *B. cereus s.l.* bacteria is limited in comparison, since these human pathogens mainly cause foodborne illness and rarely cause severe human infections. First line antibiotic treatment for human anthrax includes quinolones and tetracyclines (penicillins can be used for known sensitive isolates) [17]. For non-anthracis *B. cereus s.l.* bacteria, vancomycin (VAN) is considered a preferred first line therapy with combination or alternative treatment based on available antimicrobial susceptibility testing (AST) profiles; while penicillins are not recommended for non-anthracis *B. cereus s.l*. bacteria due to intrinsic resistance [18-20]. Understanding the AST profiles for *B. paranthracis* improves how clinicians manage infections. Identifying phenotypic variations among *B. cereus s.l*. organisms will expedite accurate *B. paranthracis* identification, can mitigate nomenclature confusion, and improve both clinical diagnosis and management.

Here, we used the same 20 clinical *B. paranthracis* strains (Table 1) from the CDC collection investigated in the Sabin et. al genetic study and perform phenotypic characterization using classical microbiological methods routinely completed for *B. anthracis* identification [21]. AST profiles for *B. paranthracis* were determined using antibiotics recommended for anthrax treatment. The suitability of a *B. anthracis* specific rapid MinION WGS sequencing workflow was also assessed for *B. paranthracis*. Our findings will help future laboratory investigations involving *B. paranthracis* or *B. anthracis*, but emphasize the need to expand detection, surveillance, and AMR testing of these pathogens.

**Table 1:**
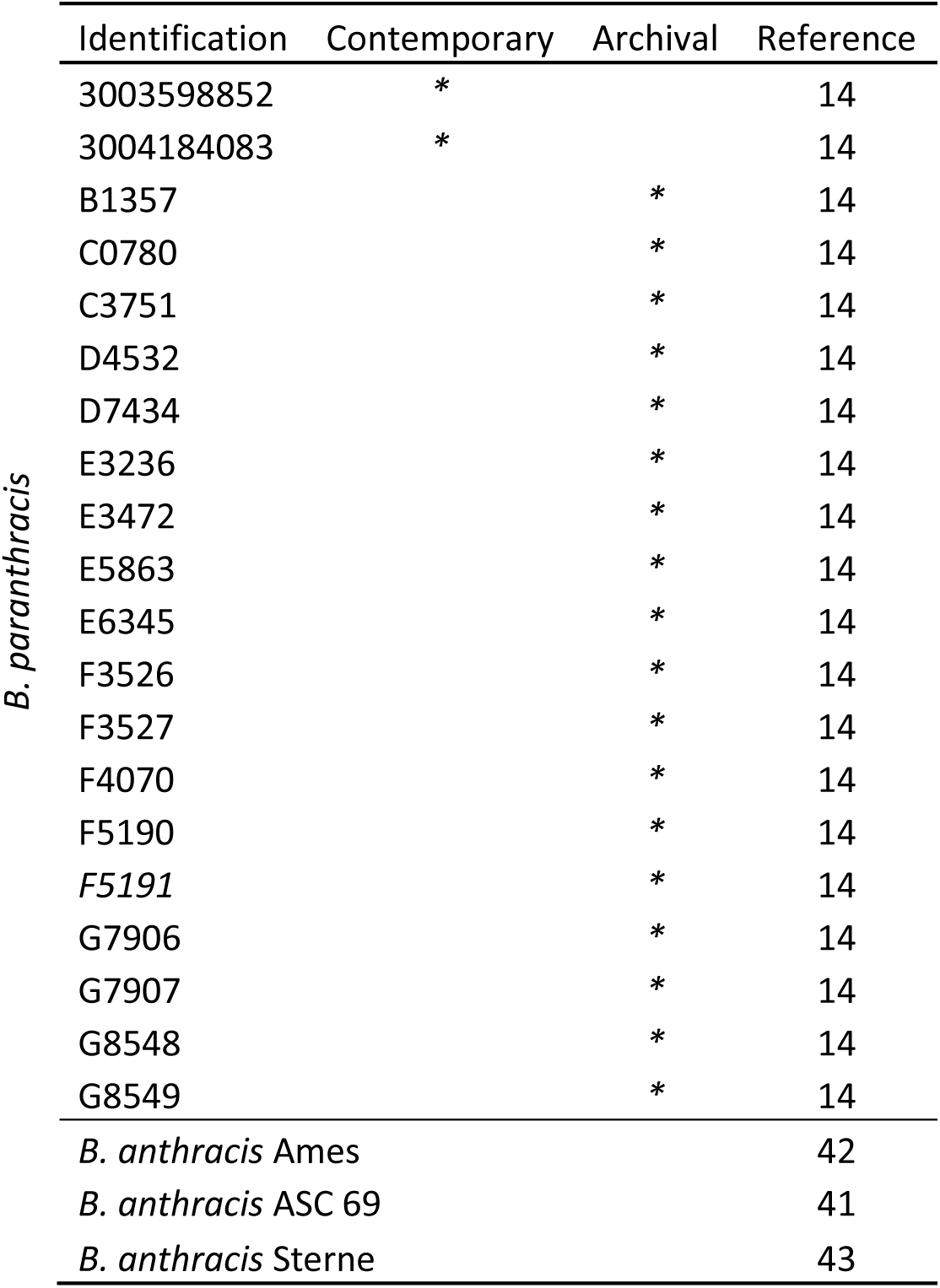
Antimicrobial Resistance Profiling and Phenotypic Characterization of Archived Clinical *Bacillus paranthracis* Strains. Michel et al 2026 Strain List

## RESULTS

### Phenotypic Characterization

Growth was observed for all 20 *B. paranthracis* strains (Table 1) on trypticase soy agar plates containing 5% sheep’s blood (SBA) after overnight incubation at 35°C in ambient air. Phenotypes for representative strains are shown in Figure 1; summary of *B. paranthracis* phenotypes compared to the expected results for *B. anthracis* rule-out testing is listed in Table 2. All strains were Gram-positive rods with rounded edges and grew in chains. Nineteen of 20 strains produced a rough ground-glass textured colony morphology on SBA (Figure 1A, and 1B), like *B. anthracis* (Figure 1D), while one strain (D7434) appeared mucoid on SBA (Figure 1C). Colony diameter ranged from 4 to 14.8 mm, appeared greyish in color, with varying morphologies between strains (e.g., round, irregular, raised, umbonate, and flat). Five strains (C3751, D7434, F5190, F5191 and G7906) were nonhemolytic on SBA. No growth was observed on MacConkey agar after 24 to 48 hours incubation at 35°C in ambient air. All strains sporulated on T3 agar when cultured in 5% CO_2_ at 35°C for 16 to 48 hours and produced spores with an oval appearance that did not swell the sporangia. (Data not shown) Spore location (central, subterminal and terminal) varied across all strains. All strains were catalase positive and oxidase negative; by semi-solid agar testing nine strains appeared non-motile like *B. anthracis* (Figure 1E and 1F) and 11 motile (Figure 1G); all strains were resistant to lysis by gamma phage (Table 2).

**Figure 1:**
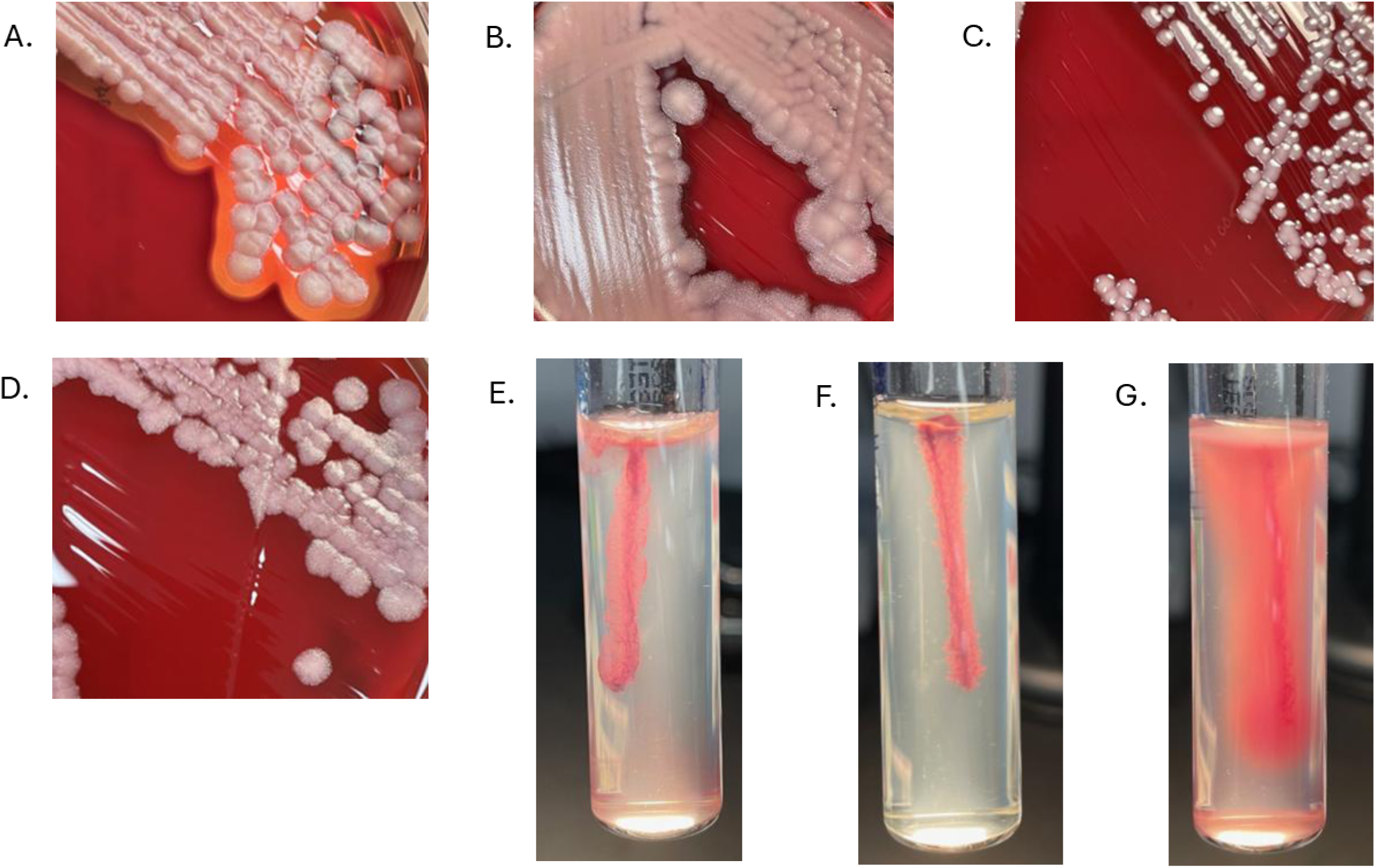
Antimicrobial Resistance Profiling and Phenotypic Characterization of Archived Clinical *Bacillus paranthracis* Strains. – Michel et al 2026 Variations of colonies characteristics in representative *B. paranthracis* strains grown on tryptic soy agar with 5% sheep’s blood (SBA) and variations of motility by semi solid agar (supplemented with TTC) testing. These observations in *B. paranthracis* were compared to characteristics of *B. anthracis*. **A.** *B. paranthracis* E3236 hemolytic and ground glass appearance on SBA. **B.** *B. paranthracis* F5191 non-hemolytic and ground glass appearance on SBA. **C.** *B. paranthracis* D7434 non-hemolytic and mucoid appearance on SBA. **D.** *B. anthracis* non-hemolytic colonies and ground glass appearance on SBA **E.** *B. anthracis*; non-motile. **F.** *B. paranthracis* C3751; non-motile. **G.** *B. paranthracis* F3527; motile.

**Table 2:**
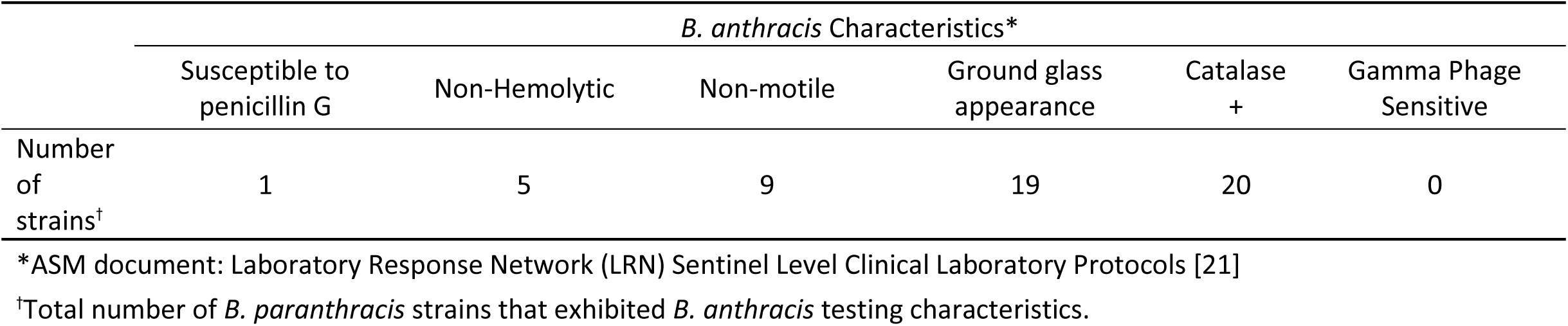
Antimicrobial Resistance Profiling and Phenotypic Characterization of Archived Clinical *Bacillus paranthracis* Strains. - Michel et al 2026 *B. paranthracis* characteristics observed across 20 CDC strains when using microbiological tests for identification of *B. anthracis*.

### Antimicrobial Susceptibility Testing

Conventional broth microdilution (BMD) testing was carried out on *B. paraanthracis* (n=20) using antibiotics recommended for treatment of anthrax disease [17]. Nineteen strains were resistant to penicillin (PEN), six strains showed elevated minimum inhibitory concentration (MIC) values to four antibiotics recommended for anthrax treatment (Table 3). Strain F5191 had an elevated MIC to ciprofloxacin (CIP). Strains E3236, F4070, and 3004184083 produced elevated MIC values to both tetracycline (TET) and doxycycline (DOX). Strains B1357 and D7434 had elevated MIC values to clindamycin (CLI). All 20 *B. paranthracis* strains were susceptible to VAN, levofloxacin (LVX), meropenem (MEM), imipenem (IPM), and antibiotics also recommended for treatment of anthrax (Table S1). Susceptibility interpretive breakpoints are not defined for some of these drugs for either *B. anthracis* or *Bacillus* spp. or are different between non-anthracis *Bacillus spp*. and *B. anthracis* [22]. For MEM and IPM, there are no defined breakpoints for *B. anthracis* and susceptibility was assigned using *Bacillus spp*. breakpoints. For LVX, the breakpoints for *B. anthracis* and *Bacillus spp*. are different (S≤0.25 and S≤2, respectively) [22]. See Supplementary Table S1 for complete AST results, susceptibility interpretive breakpoints and antibiotic testing ranges.

**Table 3:** Antimicrobial Resistance Profiling and Phenotypic Characterization of Archived Clinical *Bacillus paranthracis* Strains. - Michel et al 2026 *B. paranthracis* strains with decreased susceptibility to multiple antibiotics recommended for anthrax treatment.

| Strain |  | MIC, µg/mL |  |  |  |  |
| --- | --- | --- | --- | --- | --- | --- |
|  |  | PEN | CIP | DOX | TET | CLI |
| *B. paranthracis* | B1357 | 64 | - | - | - | 2 |
|  | D7434 | 4 | - | - | - | 1 |
|  | E3236 | 16 | - | 2 | 64 | - |
|  | F4070 | 32 | - | 2 | 32 | - |
|  | F5191 | 128 | 0.5 | - | - | - |
|  | 3004184083 | 128 | - | 4 | 64 | - |
| *B. anthracis* Ames |  | 0.12 | 0.06 | ≤0.03 | 0.06 | 0.12 |
| *B. anthracis* break points* |  | S≤0.5 | S ≤0.25 | S ≤1 | S ≤1 | - |
| *Bacillus spp.* break points* |  | S≤0.12 | S ≤1 | - | S ≤4 | S ≤0.5 |

\*Clinical & Laboratory Standards Institute M45 [22]

### Capsule Visualization

All 20 *B. paranthracis* strains were tested for capsule production by India Ink staining. Only three strains (3004184083, D7434, 3003598852) were positive (Figure 2A-C). Figure 2D depicts a *B. paranthracis* strain with no visible capsule by India Ink as a representative example of negative staining, - strain C3751. As previously reported, all three strains that were positive by India Ink, contained the genes that code for poly-y-D-glutamic acid (PGA) capsule (*capBCADE*) [14]. These three strains were further assessed for PGA presence using the InBios Active Anthrax Detect (AAD) lateral flow immunoassay (LFI), that can rapidly detect *B. anthracis* PGA [23]. Strains that were negative by India Ink staining did not contain PGA genes *capBCADE* [14], and thus were not tested by LFI. We used three culturing conditions to generate cells for the AAD-LFI assay to detect PGA in *B. paranthracis*. By this qualitative assessment, growth in heart infusion broth containing sodium bicarbonate and horse serum improved PGA visualization compared to growth in broth with bicarbonate but without horse serum. The PGA detection results from the former conditions are shown in Table 4. Strains 3004184083 and 3003598852 were weakly positive by AAD-LFI, but D7434 was negative by this test (Figure 2E). Growth on agar containing sodium bicarbonate should produce mucoid colonies for capsule producing bacteria [21]; however, for strain 3004184083, this colony phenotype was not observed despite it being positive for capsule by Inia Ink staining and for PGA by LFI (Table 4). Colonies for D7434 and 3003598852 were mucoid on sodium bicarbonate agar, agreeing with positive India Ink staining for capsule (Table 4).

**Figure 2:**
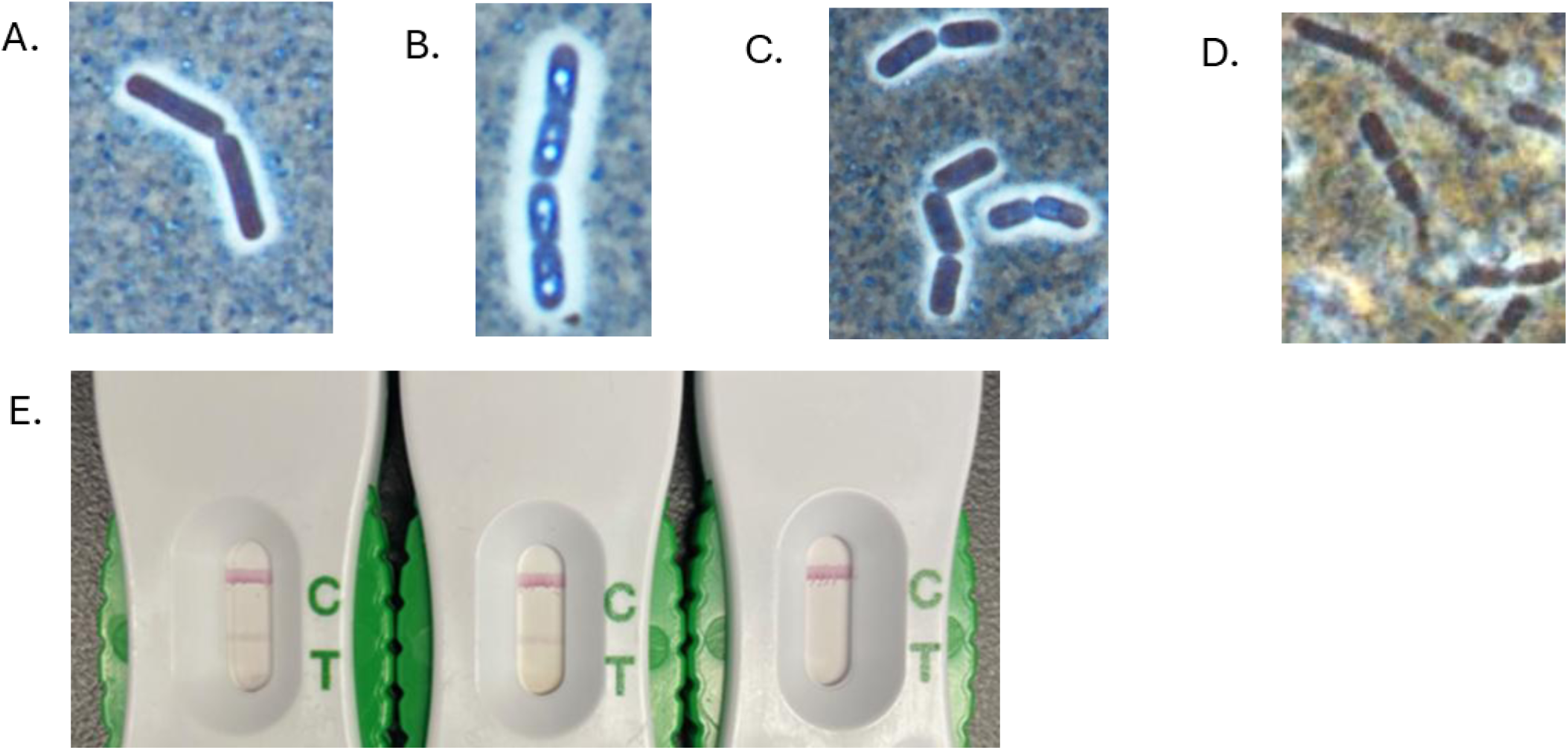
Antimicrobial Resistance Profiling and Phenotypic Characterization of Archived Clinical *Bacillus paranthracis* Strains. – Michel et al 2026 Capsule visualization by India Ink staining and detection of poly-D-glutamic acid (PGA) capsule by Active Anthrax Detect^TM^ lateral flow immunoassay (AAD-LFI) in *B. paranthracis*. **A, B,** and **C.** Positive capsule detection for strains 3004184083, D7434, and 3003598852, respectively, by India Ink staining. **D.** Negative capsule detection for G8549 by India Ink staining. **E.** The AAD-LFI cartridges show positive results for PGA presence (left, center) for 3004184083, 3003598852 and negative result for PGA (right) for D7434. ‘T’ is the Test line and ‘C’ is the Control line.

**Table 4.** Capsule detection in *B. paranthracis*.

|  | Mucoid appearance<br>(SB* Agar) | India Staining<br>(SB* agar) | AAD-Lateral Flow<br>Test (HIB+SB+HS<sup>†</sup>) | AAD-Lateral Flow<br>Test (HIB+SB<sup>‡</sup>) |
| --- | --- | --- | --- | --- |
| *B. paranthracis* 3003598852 | YES | YES | YES | YES |
| *B. paranthracis* D7434 | YES | YES | NO | NO |
| *B. paranthracis* 3004184083 | NO | YES | YES | YES |
| *B. anthracis* ASC 69 | YES | YES | YES | YES |

\*Sodium Bicarbonate
<sup>†</sup>Heart Infusion Broth + Sodium Bicarbonate + Horse Serum
<sup>‡</sup>Heart Infusion Broth + Sodium Bicarbonate

### Same-day Whole Genome Sequencing Analysis

Following the same-day MinION sequencing workflow used previously for a clinical strain of *B. anthracis* [24], genomic DNA prepared from each of 20 *B. paranthracis* strains met the minimal quality requirements for sequencing, but failed to produce complete chromosome and plasmid assemblies using the PIMA 1.0 software. *B. paranthracis* sequencing produced open contigs and, for the drug-resistant strains, did not correctly detect AMR genes (Supplementary Table S2).

## DISCUSSION

*B. paranthracis* was proposed as a novel species based on polyphasic taxonomy, however due to nomenclature similarity it has been confused with *B. anthracis* resulting in delayed responses to outbreak investigations [1, 3]. It is of critical importance in clinical and public health contexts to establish accurate identification of *B. paranthracis* and other non-*anthracis B. cereus s.l.* strains and distinguish them from *B. anthracis* using standard microbiological methods. Non-foodborne human infections with non-*anthracis B. cereus s.l*. appear to be rare. To understand the frequency of such infections, Sabin et al. investigated diversity within the CDC collection of non-*anthracis B. cereus s.l*. strains from non-foodborne illness [14]. For these archival (1967-2003) and contemporary (2019-2023) non-*anthracis B. cereus s.l*. strains, there were associated reports of human infections with *B. paranthracis* which included descriptions of “sepsis”, “fatal sepsis”, “skin lesions”, and “pneumonia” [14]. Additionally, a recent study identified *B. paranthracis* as the predominant species associated with invasive infections in preterm neonates across 14 French hospitals [25]. These are serious infections, therefore correct diagnosis of *B. paranthracis* would help select appropriate treatment and remove the regulatory burden associated with reporting a select agent such as *B. anthracis*.

For this investigation, 20 *B. paranthracis* strains (not associated with foodborne illness) were selected from the CDC collection for further testing beyond the genetic characterization previously reported [15]. The classical microbiological methods used for identification of *B. anthracis* do not offer a consistent profile that is useful for *B. paranthracis* identification. Unlike *B. anthracis*, hemolysis on SBA, motility and colony morphology varied across *B. paranthracis* strains. Like *B. anthracis,* all *B. paranthracis* strains were catalase positive, oxidase negative and did not grow on McConkey agar. Every *B. paranthracis* strain in this study was resistant to gamma phage lysis and this was the single consistent phenotype different from *B. anthracis*, but this feature is also shared by other non-anthracis *B. cereus s.l.* species [26, 27]. Lysis by gamma phage is described as 97 and 96% specific for *B. anthracis* when used as diagnostic assays [25, 26]. *B. paranthracis* is taxonomically similar to *B. anthracis* and with the exceptions of gamma phage sensitivity for 20 strains assessed in this study, there are not many reliable phenotypic distinctions between them.

PGA capsule is a virulence factor that helps *B. anthracis* evade the host immune system during infection, and detection of PGA has been used as part of *B. anthracis* identification [28]. The presence of the anthrax PGA capsule gene operon *capBCADE* in some of *B. paranthracis* strains, reported previously [14], demonstrated how reliance on these targets can confound accurate discrimination of *B. paranthracis* from *B. anthracis.* When *B. paranthracis* strains were subjected to India Ink staining, D7434 and two additional strains, which all carry capsule genes *capBCADE,* were positive for capsule presence. Surprisingly, *B. paranthracis* strain D7434 was negative for PGA when using the AAD-LFI assay. Possible explanations for negative AAD-LFI results in D7434 are: *capBCADE* operon is expressed but gene products are modified and are not recognized by this test, or *capBCADE* is not expressed and D7434 makes a non-PGA capsule. Genome analysis from a recent publication demonstrated presence of genes for a polysaccharide capsule in *B. paranthracis* [12], which is distinct from a PGA capsule. Polysaccharide capsules are common in the *B. cereus* group and other bacterial pathogens. This suggests that non-PGA capsules might be expressed in *B. paranthracis*. In this study, we did not investigate this further and did not analyze genome sequences for capsule genes since we were focused on phenotypic testing. Specific culturing conditions for detection of capsule in *B. anthracis* and other *B. cereus s.l.* organisms described in previous studies were mostly useful for capsule identification in *B. paranthracis* [27, 29-32]. Capsule detection by India ink staining, mucoid colony phenotype on bicarb agar and detection of PGA using the AAD-LFI assay are methods used for identification of *B. anthracis*. However, this is not appropriate for identification of *B. paranthracis*, as most strains were negative for capsule production, and a strain positive for capsule production failed to produce a mucoid colony phenotype on bicarb as previously reported in *B. cereus s.l.* [27].

AST profiles with resistance to multiple drug classes were identified in *B. paranthracis* strains that have not been observed in *B. anthracis*. All *B. paranthracis* strains, except one, were resistant to PEN, amoxicillin (AMX), amoxicillin-clavulanic acid (AMC), and ampicillin (AMP). Resistance to these antibiotics is common in *B. cereus s.l.* bacteria and susceptibility to PEN is historically a method used in identification of *B. anthracis* [28]. With a single PEN susceptible strain, PEN-susceptibility may not be reliable to distinguish *B. paranthracis* and *B. anthracis*. Six *B. paranthracis* strains showed elevated MIC values to four antibiotics recommended for treatment of anthrax: CIP, TET, DOX, and clindamycin (CLI). CIP and CLI are considered first-line and alternative antibiotics for anthrax treatment, and in *B. anthracis*, MICs of 0.25 mg/ml CIP and 0.12 mg/ml CLI are the maximum values reported in wild type strains [17, 33]. Resistance to TET (MIC >16 by Sensititre AST panel), an MIC of 0.5 for CIP (Sensititre AST panel) and resistance to MIN by Etest in the *B. paranthracis* were observed in the Mn5 type strain (data not shown). Currently, only a single published report exists of naturally occurring TET resistance in *B. anthracis* [33]; here, we identified three *B. paranthracis* strains that are resistant to TET and DOX. Like most *B. cereus s.l.* bacteria, all *B. paranthracis* strains were susceptible to VAN. Moreover, the susceptibility profiles of *B. paranthracis* strains in this investigation are similar to the AST profiles previously reported for *B. cereus s.l.* strains where authors describe few strains with resistance to clinically important drugs such as LVX, CLI and linezolid (LZD) that have led to positive clinical outcomes [18, 34-36]. For *Bacillus spp*., CLSI recommends primary susceptibility testing for VAN, fluoroquinolones, and CLI, but includes breakpoints for VAN, PEN, AMP, IPM, MEM, gentamycin, amikacin, erythromycin, CIP, levofloxacin, CLI, sulfamethoxazole/trimethoprim, chloramphenicol, and rifampin [22]. Since VAN is recommended for treatment of *B. cereus* infections and clinical strains appear susceptible, surveillance and expanded susceptibility profiling of additional *B. paranthracis* strains are warranted to offer a more complete picture [18, 19]. Based on our AST results, following guidelines for empirical therapy for *B. cereus* would be recommended for *B. paranthracis* infection treatment [18, 36].

In a recent report*, B. paranthracis* was isolated from immunocompromised infants with bacteremia and has been reported to be the causative agent of sepsis in a 26-year-old patient with T-cell acute lymphoblastic leukemia [12, 37]. Suzuk Yildiz, et al. reported the *B. paranthracis* isolated from the T-cell acute lymphoblastic leukemia patient as IPM-, MEM-, and VAN-susceptible, and also described as susceptible to CIP and levofloxacin “in a dose dependent manner” by disc diffusion [12]. Susceptible-Dose Dependent (SDD) or susceptible increased exposure (I) is a category for AST introduced by the European Committee on Antimicrobial Susceptibility Testing (EUCAST) in 2019 [38]. The AST profiles of strains in this category depend on the dosing regimens used, where higher doses or more frequent doses or both are used to achieve a concentration level that is likely to be clinically effective against the strain while giving proper consideration to the maximum approved dosage regimen [38]. In this study, we identified a single *B. paranthracis* strain considered “non-susceptible” to CIP based on CLSI MIC interpretive criteria for *B. anthracis,* but “susceptible” to CIP based on CLSI breakpoints for *Bacillus spp.* by BMD. Considering the potential overlap in disease presentation for some *B. paranthracis* infections with anthrax, the challenges of interpreting *in vitro* AST results can complicate decisions about treatment.

New, rapid AST profiling methods have been developed for different pathogens, but not *B. paranthracis* [24]. In 2020, MinION sequencing and bioinformatic analysis using PiMA v1.0 was used to rapidly characterize *B. anthracis* during an outbreak investigation including detection of known antimicrobial resistance (AMR) markers [39]. This method correctly ruled out evidence of AMR genes or introduced plasmid(s) within hours of receiving a human anthrax strain [39]. We found that the published same-day MinION sequencing procedure and custom bioinformatics pipeline developed for *B. anthracis* was not suitable with *B. paranthracis*. PiMA v1.0 software was developed to rapidly characterize *B. anthracis* genomes using nanopore sequencing, relying on a well-defined reference genome and a highly curated list of AMR gene mutations. It is optimized to detect plasmids, structural integrations, and resistance markers rapidly [24], and performs well with *B. anthracis* because of its clonal genome structure and low variability. The rapid sequencing procedure utilizes gDNA extracted directly from an overnight culture grown on SBA and the approach was applied for *B. paranthracis*. However, with *B. paranthracis*, this approach did not produce reliable results with downstream analysis using PiMA v1.0. With ∼95% ANI with *B. anthracis*, and more genome variability [1, 14], the *B. paranthracis* assemblies were often fragmented. As a result, PiMA v1.0 reported a high number of variants, most of which likely represent species level differences rather than true resistance mutations. This limits the accuracy and usefulness of PiMA v1.0 for rapid genomic analysis in non-*B. anthracis* strains, unless it is adapted to include a robust species-specific reference and relevant AMR mutations. For additional information regarding adaptation of PiMA for genomic analysis of *B. paranthracis*, (Chivukula et al., in preparation).

Our investigation is limited by the low number of *B. paranthracis* strains included (n=20). With few reports of *B. paranthracis* infection, the clinical significance might be incomplete and potentially due to unfamiliarity with this species. Additionally, the susceptibility interpretations described in this study rely on *B. anthracis* and *Bacillus spp*. breakpoints which might not be appropriate for a different species.

Here we report that the taxonomically similar *B. paranthracis* and *B. anthracis* species are diverse when assessing microbiological phenotypes and AST profiles. Expanding surveillance and characterization of additional *B. paranthracis* isolates will strengthen our ability to accurately identify and manage *B. paranthracis* infections. The continued genomic characterization of *B. paranthracis* strains, laboratory based microbiological characterization, AST profiling, and AMR identification are critical to future public health investigations and for treatment guidance.

## MATERIALS AND METHODS

### Biosafety Procedures

All procedures with bacterial select agents were performed in a biosafety level 3 (BSL-3) laboratory by trained personnel wearing appropriate personal protective equipment (PPE). All procedures with select agent excluded bacterial strains were performed in a biosafety level 2 (BSL-2) laboratory by trained personnel wearing appropriate PPE. All manipulations of cell cultures were performed in a class II type A2 biological safety cabinet as recommended by the CDC/NIH publication Biosafety in Microbiological and Biomedical Laboratories, 6th edition [40].

### Bacterial Strains

All *B. paranthracis* strains were archived at -70°C in brain heart infusion broth containing 20% glycerol and streaked as pure cultures on SBA (Remel, R01202) for this study. *B. anthracis* strains (Sterne, Ames, ASC 69) were used as controls throughout this study [41-43].

### Phenotypic Characterization

Strains were streaked on to SBA and grown at 35°C in ambient air for 16-20 hours to observe hemolysis and colony morphologies, measure colony sizes, and perform oxidase and catalase testing. Strains were streaked on MacConkey agar and incubated at 35°C for 16-20 hours in ambient air, then observed for growth. Cellular motility was assessed using semi-solid agar containing tetrazolium chloride (TTC) (Remel, R061410). Sporulation was induced as previously described using T3 agar [44]. A Nikon Eclipse Ni-U phase contrast microscope with 100x oil emersion objective was used to observe cellular morphologies and sporulation. India Ink staining was used for capsule visualization as previously described [27]. Images of India ink stains were captured using the Nikon DS-L3 CCD microscope camera.

To assess the capsule bond to cellular surface, cells from bicarbonate agar plates were heated at 60°C for 30 min, cooled, and stained with India ink as previously described [27]. The AAD-LFI (InBios, Washington, USA) was used for detection of the capsular PGA following the manufacturer’s guidance. Three culturing conditions were assessed in conjunction with the AAD LFI: 1) Bicarbonate agar 2) Heart infusion broth containing 0.8% sodium bicarbonate and 3) Heart infusion broth supplemented with horse serum (Fisher, SR0035C) and 0.8% sodium bicarbonate. *B. paranthracis* colonies grown on bicarbonate agar were assessed for mucoid appearance as an indicator of capsule production [21]. *B. anthracis* ASC 69 [43] was used as positive control for the InBios AAD-LFI assay, India Ink staining, and as an example of mucoid appearance for colonies grown on bicarbonate agar.

### Antimicrobial Susceptibility Testing

BMD testing was conducted in accordance with the CLSI guidelines [22]. Cells from four to six isolated colonies from an overnight SBA culture were suspended in saline solution to achieve a turbidity equivalent to a 0.5 McFarland standard. Each suspension was diluted at a ratio of 1:20 in additional saline. BMD AST panels, prepared in-house with cation-adjusted Mueller-Hinton broth, were inoculated with cell suspensions to result in additional 1:10 dilution, then incubated at 35°C in ambient air for 16 to 20 hours. Antibiotics included in AST panels: PEN, AMX, AMC, AMP, IPM, MEM, CLI, clarithromycin (CLR), LZD, DOX, TET, minocycline (MIN), CIP, LVX, moxifloxacin (MXF), and VAN. The MIC of each antibiotic was recorded as concentration at the first well exhibiting no visible growth.

### Genomic DNA Extraction and Same-day Whole Genome Sequencing Analysis

Genomic DNA (gDNA) was extracted, and nanopore sequencing libraries were prepared (Rapid Barcoding Sequencing Kit SQK-RBK114.24; Oxford Nanopore Technologies) and analyzed by PiMA v1.0 as previously reported [39]. Extracted gDNA was assessed for quantity and quality using the Qubit dsDNA BR Assay Kit and NanoDrop 2000 spectrophotometer following manufacturer’s instructions.

## Acknowledgements

We would like to thank Dr. Christopher Hsu and Dr. Melisa Shah for kindly providing their expertise and experience in support of this investigation.

## Funding

This work was supported by the U.S. CDC Office Readiness and Response and the CDC Office of Advanced Molecular Detection.

The findings and conclusions in this report are those of the authors and do not necessarily represent the official position of the U.S. Centers for Disease Control and Prevention. The mention of company names or products does not constitute endorsement by CDC.

## Notes

### Competing Interest Statement

The authors have declared no competing interest.

