## Supplemental Table S1 for "Antimicrobial Resistance Profiling and Phenotypic Characterization of Archived Clinical *Bacillus paranthracis* Strains"

**Supplementary Table S1.**

| | | | | | | | | | | | | | | | | |
|---|---|---|---|---|---|---|---|---|---|---|---|---|---|---|---|---|
| *B. anthracis* susceptible brkpts (CLSI M45) | S≤0.5 | S ≤0.12 | - | - | - | - | - | - | - | S ≤1 | S ≤1 | - | S ≤0.25 | S ≤0.25 | - | - |
| *Bacillus* spp. susceptible brkpts (CLSI M45) | S ≤0.12 | - | - | S ≤0.25 | S ≤4 | S ≤4 | S ≤0.5 | - | - | - | S ≤4 | - | S ≤1 | S ≤2 | - | S ≤4 |
| | **Antibiotic** | | | | | | | | | | | | | | | |
| **Strain** | **PEN** | **AMX** | **AMC** | **AMP** | **IPM** | **MEM** | **CLI** | **CLR** | **LZD** | **DOX** | **TET** | **MIN** | **CIP** | **LVX** | **MXF** | **VAN** |
| **3004184083** | 128 | 64 | 8/4 | 32 | ≤0.03 | ≤0.03 | 0.5 | 0.5 | 2 | 4 | 64 | 2 | 0.25 | 0.25 | 0.25 | 1 |
| **3003598852** | 2 | 2 | 0.25/0.5 | 0.5 | ≤0.03 | ≤0.03 | 0.5 | 0.5 | 2 | 0.12 | 0.5 | 1 | 0.25 | 0.12 | 0.12 | 1 |
| **B1357** | 64 | 32 | 4/2 | 16 | ≤0.03 | 0.06 | 2 | 0.5 | 2 | 0.06 | 0.5 | 0.5 | 0.25 | 0.25 | 0.25 | 1 |
| **C0780** | 4 | 2 | 1/0.5 | 2 | ≤0.03 | 0.06 | 0.5 | 0.12 | 2 | ≤0.03 | 0.06 | ≤0.03 | 0.25 | 0.25 | 0.25 | 1 |
| **C3751** | 0.03 | 0.03 | 0.03/0.015 | 0.06 | ≤0.03 | 0.06 | 0.5 | 0.25 | 2 | 0.06 | 1 | 0.5 | 0.12 | 0.12 | 0.12 | 1 |
| **D4532** | 16 | 4 | 2/1 | 4 | ≤0.03 | 0.06 | 0.5 | 0.12 | 1 | 0.12 | 0.5 | 0.5 | 0.25 | 0.25 | 0.12 | 1 |
| **D7434** | 4 | 2 | 1/0.5 | 2 | ≤0.03 | ≤0.03 | 1 | 0.25 | 2 | 0.25 | 1 | 1 | 0.25 | 0.25 | 0.12 | 1 |
| **E3236** | 16 | 8 | 4/2 | 16 | 0.5 | ≤0.03 | 0.5 | 0.12 | 2 | 2 | 64 | 0.5 | 0.12 | 0.12 | 0.12 | 1 |
| **E3472** | 4 | 4 | 2/1 | 4 | ≤0.03 | ≤0.03 | 0.5 | 0.12 | 2 | 0.06 | 0.25 | 0.25 | 0.25 | 0.12 | 0.12 | 1 |
| **E5863** | 8 | 4 | 2/1 | 8 | ≤0.03 | 0.06 | 0.25 | 0.25 | 2 | ≤0.03 | 0.06 | ≤0.03 | 0.25 | 0.25 | 0.12 | 1 |
| **E6345** | 2 | 0.5 | 0.5/0.25 | 1 | ≤0.03 | ≤0.03 | 0.5 | 0.25 | 2 | 0.06 | 0.5 | 0.25 | 0.12 | 0.12 | 0.12 | 1 |
| **F3526** | 4 | >2 | 2/1 | 4 | 0.03 | 0.03 | 0.5 | 0.12 | 1 | 0.12 | 0.5 | 0.5 | 0.12 | 0.12 | 0.12 | 1 |
| **F3527** | 4 | 4 | 2/1 | 4 | ≤0.03 | ≤0.03 | 0.5 | 0.12 | 2 | 0.12 | 0.5 | 0.25 | 0.12 | 0.12 | 0.12 | 1 |
| **F4070** | 32 | 8 | 2/1 | 16 | ≤0.03 | ≤0.03 | 0.5 | 0.5 | 1 | 2 | 32 | 0.5 | 0.06 | 0.06 | 0.06 | 1 |
| **F5190** | 4 | 2 | 1/0.5 | 2 | ≤0.03 | ≤0.03 | 0.5 | 0.25 | 2 | 0.12 | 0.5 | 0.5 | 0.06 | 0.06 | 0.03 | 1 |
| **F5191** | 128 | 32 | 4/2 | 64 | ≤0.03 | 0.06 | 0.5 | 0.25 | 2 | 0.06 | 0.5 | 0.5 | 0.5 | 0.25 | 0.25 | 1 |
| **G7906** | 2 | 4 | 2/1 | 4 | 0.25 | 0.06 | 0.5 | 0.25 | 2 | ≤0.03 | 0.06 | 0.06 | 0.25 | 0.25 | 0.25 | 1 |
| **G7907** | 8 | 8 | 4/2 | 8 | ≤0.03 | ≤0.03 | 0.5 | 0.25 | 2 | 0.06 | 0.5 | 0.5 | 0.12 | 0.12 | 0.12 | 1 |
| **G8548** | 4 | 2 | 2/1 | 4 | ≤0.03 | ≤0.03 | 0.5 | 0.12 | 2 | 0.12 | 0.5 | 0.5 | 0.12 | 0.12 | 0.12 | 1 |
| **G8549** | 4 | 4 | 4/2 | 4 | ≤0.03 | ≤0.03 | 0.5 | 0.25 | 2 | 0.06 | 0.25 | 0.25 | 0.12 | 0.12 | 0.12 | 1 |
