## Supplemental Table S2 for "Antimicrobial Resistance Profiling and Phenotypic Characterization of Archived Clinical *Bacillus paranthracis* Strains"

Table S2. Summary statistics of genome assembly outputs and similarity to *Bacillus anthracis* generated by PiMA.

| strain | genome size (bp) | number contigs | closed chromosome? | number open contigs | average chromosome coverage (X) | ANI (%)* | Coverage (%)** | n_amr_gene | n_amr_snp |
|---|---|---|---|---|---|---|---|---|---|
| C3751 | 5882545 | 10 | Y | 4 | 177 | 95.4 | 81.28 | 0 | 423 |
| E6345 | 5293710 | 4 | Y | 1 | 183 | 95.3 | 81.59 | 0 | 343 |
| G7906 | 6253423 | 8 | Y | 2 | 166 | 95.3 | 82.89 | 0 | 336 |
| F5191 | 5254757 | 3 | Y | 1 | 207 | 95.5 | 82.1 | 0 | 322 |
| F3526 | 5760905 | 7 | Y | 1 | 179 | 95.2 | 83.59 | 0 | 343 |
| B1357 | 5464127 | 5 | Y | 2 | 200 | 95.5 | 82.61 | 0 | 411 |
| E3236 | 5837728 | 8 | Y | 3 | 176 | 95.2 | 83.6 | 0 | 343 |
| G8548 | 5637509 | 7 | Y | 3 | 186 | 95.2 | 83.04 | 0 | 343 |
| F3527 | 5739502 | 7 | Y | 1 | 183 | 95.2 | 83.59 | 0 | 343 |
| G7907 | 5771203 | 8 | Y | 2 | 184 | 95.2 | 83.52 | 0 | 343 |
| D7434 | 5488990 | 8 | Y | 4 | 177 | 95.5 | 82.79 | 0 | 453 |
| G8549 | 5630465 | 11 | Y | 5 | 177 | 95.2 | 83.05 | 0 | 343 |
| C0780 | 5372618 | 12 | Y | 7 | 189 | 95.5 | 82.58 | 0 | 430 |
| E3472 | 5753213 | 15 | Y | 11 | 183 | 95.2 | 83.58 | 0 | 342 |
| D4532 | 5644411 | 16 | N | 12 | 58 | 95.3 | 84.21 | 0 | 410 |
| 3004184083 | 5559795 | 16 | N | 13 | 194 | 95.4 | 84.37 | 1 | 417 |
| E5863 | 5644230 | 21 | N | 15 | 128 | 95.2 | 82.02 | 0 | 325 |
| F4070 | 5867293 | 22 | Y | 14 | 153 | 95 | 84.1 | 0 | 382 |
| F5190 | 5365923 | 28 | N | 22 | 101 | 95.4 | 79.89 | 0 | 411 |
| 3003598852 | 5917032 | 44 | N | 28 | 118 | 95.4 | 83.03 | 0 | 427 |

* Average Nucleotide Identity - calculated from homologous regions from the assembled genome and the reference

** Percent alignment of the assembly with the reference genome
